# Establishment of multiple novel patient-derived models of desmoplastic small round cell tumor enabling functional characterization of ERBB pathway signaling and pre-clinical evaluation of a novel targeted therapy approach

**DOI:** 10.1101/2020.09.22.308940

**Authors:** Roger S. Smith, Igor Odintsov, Zebing Liu, Allan Jo-Weng Lui, Takuo Hayashi, Morana Vojnic, Yoshiyuki Suehara, Lukas Delasos, Marissa S. Mattar, Julija Hmeljak, Hillary A. Ramirez, Melissa Shaw, Gabrielle Bui, Alifiani B. Hartono, Eric Gladstone, Siddharth Kunte, Heather Magnan, Inna Khodos, Elisa De Stanchina, Michael P. La Quaglia, Jinjuan Yao, Marick Laé, Sean B. Lee, Lee Spraggon, Christine A. Pratilas, Marc Ladanyi, Romel Somwar

**Affiliations:** Department of Pathology, Memorial Sloan Kettering Cancer Center, New York, NY, USA; Human Oncology and Pathogenesis Program, Memorial Sloan Kettering Cancer Center, New York, NY, USA; Anti-tumor Assessment Core Facility, Molecular Pharmacology Program, Memorial Sloan Kettering Cancer Center, New York, NY, USA; Gerstner School of Graduate Studies, Memorial Sloan Kettering Cancer Center, New York, NY, USA; Tulane University School of Medicine, New Orleans, LA, USA; Department of Pediatrics, Memorial Sloan Kettering Cancer Center, New York, NY, USA; Department of Surgery, Memorial Sloan Kettering Cancer Center, New York, NY, USA; Division of Pediatric Oncology, Sidney Kimmel Comprehensive Cancer Center at Johns Hopkins, Baltimore, MD, USA

**Keywords:** EWSR1-WT1, DSRCT PDX, EGFR, sarcoma proteomics

## Abstract

Desmoplastic small round cell tumor (DSRCT) is characterized by the t(11;22)(p13;q12) chromosomal translocation, which fuses the transcriptional regulatory domain of *EWSR1* with the zinc finger DNA-binding domain of *WT1*, resulting in the oncogenic transcription factor EWS-WT1. DSRCT primarily affects young males and has a 5-year overall survival of about 15%. Typical treatment approaches for patients with DSRCT involve a multi-modal combination of surgery, chemotherapy and radiation. The paucity of DSRCT disease models has hampered functional and pre-clinical therapeutic studies in this aggressive cancer. Here, we developed robust preclinical disease models and mined DSRCT expression profiling data to identify genetic vulnerabilities that could be leveraged for the identification of rational therapies. Specifically, we developed four new DSRCT cell lines and one patient-derived xenograft (PDX) model. Transcriptomic and proteomic profiling showed evidence of activation of the ERBB pathway. Ectopic expression of EWSR1-WT1 resulted in upregulation of ERRB family ligands and downstream signaling. Treatment of DSRCT cell lines with ERBB ligands resulted in activation of EGFR, ERBB2, ERK1/2 and AKT, and stimulation of cell growth. Conversely, targeting of EGFR using shRNA, small molecule inhibitors (afatinib, neratinib) or an anti-EGFR antibody (cetuximab) inhibited growth and induced apoptosis in DSRCT cells. Finally, treatment of mice bearing DSRCT xenografts with a combination of cetuximab and afatinib significantly reduced tumor growth. These data provide a rationale for the clinical evaluation of EGFR antagonists in patients with DSRCT.

## Introduction

Desmoplastic small round cell tumor (DSRCT) is a devastating sarcoma of primitive histology that affects pediatric and adolescent patients (1). Tumors are characterized by the recurrent chromosomal translocation t(11;22)(p13;q12) (2,3) which results in the fusion of the first 7, 9, or 10 exons of *EWSR1*, to exon 8 of *WT1*, coupling the transcriptional regulatory domain of *EWSR1* and the DNA-binding domain of *WT1* (4–6). Patients with DSRCT have poor overall outcomes despite initial responses to aggressive treatments, including surgery, chemotherapy and radiation (7). Complete surgical resection is challenging due to the abundance of disseminated lesions at multiple peritoneal sites at the time of diagnosis (7). In a report on the longitudinal experience over 30 years with a cohort of 66 patients, overall survival was a dismal 15% at five years (8).

The development of targeted therapy for DSRCT and other sarcomas driven by chimeric transcription regulators has lagged behind that for other solid tumors. One of the main reasons is that targeting the primary drivers with small molecules that need to penetrate the nuclear compartment is challenging due to the non-enzymatic nature of chimeric transcription regulators and the complexity of their binding to DNA. To improve clinical outcomes for this aggressive adolescent and young adult sarcoma, it will be necessary to identify alternative therapeutic strategies.

Protein kinases may represent genetic vulnerabilities that can be exploited as potential therapeutic targets for DSRCT. As kinase signaling is central to cell proliferation and survival, it is likely that these aberrant transcription factors achieve their effects on cell growth through the dysregulation of specific kinase signaling pathways. Identifying these upregulated kinases, which remain poorly defined for DSRCT and other sarcomas, may highlight therapeutic vulnerabilities that are more readily targetable than the fusion protein themselves. We have previously demonstrated the value of this approach in another translocation-positive sarcoma, alveolar soft part sarcoma (ASPS), where the ASPL-TFE3 fusion upregulates the MET receptor tyrosine kinase, creating a targetable dependency on MET signaling (9,10). In addition, we have previously demonstrated the sensitivity of a subset of synovial sarcoma (SS) to inhibition of PDGFRA, a kinase often upregulated in this disease (11). In Ewing sarcoma (ES) and SS, some patients have been observed to respond to IGF1R inhibitors and pazopanib (a multi-targeted receptor tyrosine kinase inhibitor), respectively (12–15), indicating targetable dependencies on kinase signaling in these two sarcomas, though the precise mechanisms of those dependencies have remained elusive.

We hypothesize that similar to SS, ES and ASPS, DSRCT has unique dependencies on kinases that can be exploited for therapy. Although DSRCT and the oncogenic driver transcription factor were identified approximately 30 years ago, studying the biology of this tumor to identify potential therapeutic targets has been limited by the paucity of preclinical disease models. Indeed, there is only one published, widely available DSRCT cell line, JN-DSRCT-1, first described in 2002 (16). The development of effective targeted therapy requires disease models such as isogenic and patient-derived cell lines and xenograft models that maintain their original characteristics and are easily manipulated for molecular genetic approaches and drug development studies. Our goal in this study was to generate new patient-derived DSRCT cell lines and xenograft models and use these to identify kinases that may serve as therapeutic targets. We report here four new fully characterized cell lines and one PDX model and show that the ERBB pathway is essential for growth of DSRCT cells and can be exploited for therapy. Importantly, we report the first orthotopic xenograft tumor models of DSRCT and demonstrate significant reduction in tumor burden with EGFR-targeted therapies.

## Materials and Methods

Cell culture media, antibiotics and phosphate-buffered saline (PBS) were prepared by the MSK Media Preparation Core Facility. Fetal bovine serum (FBS) was procured from Atlanta Biologicals (Flowery Branch, GA). HEK-293T cells were obtained from American Type Culture Collection (Manassas, VA). The JN-DSRCT1 cell line was a gift from Dr. Jun Nishio (Fukuoka University, Fukuoka, Japan). The LP-9 cell line is an untransformed, diploid, mesothelial cell line that was derived from a 26-year old female ovarian cancer patient (17) and was obtained from the Coriell Institute For Medical Research (Camden, NJ). Antibodies to all native and/or phosphorylated proteins (see Supplementary Table S1 for details) except for WT1 were purchased from Cell Signaling Technology (Boston, MA). The C-terminal anti-WT1 antibody (C-19) was purchased from Santa Cruz Biotechnology (Dallas, TX). Small molecule inhibitors were obtained from Selleckchem (Houston, TX). Promega’s ApoOne Homogenous Caspase 3/7 activity assay kit, AlamarBlue, puromycin, geneticin, tissue culture plastic wares and all Western blotting reagents were obtained from ThermoFisher Scientific (Waltman, MA). Protease inhibitor cocktail, RIPA buffer (10X) and all other chemicals not listed above were purchased from EMD-Millipore Sigma (St. Louis, MO). All oligonucleotides used for PCR assays were obtained from Integrated DNA technologies (Coralville, IA).

### Generation and growth of cell lines

Tissue samples were collected under an institutional IRB-approved biospecimen collection protocol and informed consent was obtained from patients. The SK-DSRCT-1 cell line was generated from ascites fluid. Briefly, ascites fluid was centrifuged at 1,000 RPM for 5 min to pellet cells and then pellets were washed twice with icecold PBS, centrifuged again and finally resuspended in DMEM:F12 (1:1 ratio) growth medium supplemented with 10% FBS and 1% antibiotics. The cell lines were considered established after growing for 20 continuous passages. The SK-DSRCT2 cell line was created from a surgically resected abdominal tumor which was cut into small pieces with a scalpel in serum-free DMEM:F12 growth media and then digested for 1 h with collagenase (2 mg/mL) in a final volume of 5 mL, at 37°C. The sample was vortexed every 5 mins and then DME:F12+10% FBS media was added to a final volume of 50 mL, centrifuged to pellet cells and then plated in DMEM:F12 growth medium supplemented with 10% FBS and 1% antibiotics. The sample was allowed to propagate over multiple generations, trypsinized when necessary to subculture and eventually only single cells remained. Unless indicated otherwise, all cell lines were maintained in DMEM:F12 growth medium supplemented with 10% FBS and 1% antibiotics for experiments in a humidified incubator infused with 5% CO_2_ and sub-cultured when stock flasks reached approximately 75% confluence at a 1:3 dilution. BER-DSRCT and BOD-DSRCT cell lines were derived from DSRCT patient-derived xenograft tissues as described previously (18). For determination of doubling time, cells were plated at a density of 25,000 (JN-DSRCT-1, SK-DSRCT1, BOD-DSRCT, BER-DSRCT) or 50,000 (SK-DSRCT2) cells per well in 6-well plates and then counted every 24 h. Data points were fitted to an exponential growth equation using GraphPad Prism to determine the doubling time. Clonogenic assays were used to determined the plating efficiency of each cell line (19). Cells were plated at 3 densities in 6 mm dishes (1,000, 2,000 or 3,000 cells per dish), in triplicates and grown for two weeks. Colonies were then stained with crystal violet and counted. Plating efficiency was determined as the # of colonies/# of cells plated for each density and then averaged. Only colonies of >100 cells were counted. For ligand-induced growth, BER-DSRCT and SK-DSRCT2 cells were plated in 12-well plates at a density of 10,000 cells/well and 24 h after plating (day 0) cells were counted and then placed in media containing 10% FBS, 1% FBS or 1% FBS supplemented with 100 ng/mL EGF or HB-EGF. Cells were counted every 24-48 h thereafter. For immunoprecipitation studies, cells were plated a density of 5 million cells in 10-cm dishes. Two days later, cells were depleted of serum for 24 h before preparation of cell extracts.

### Spectral and DAPI Karyotyping

Metaphase preparations were hybridized with spectral karyotyping (SKY) painting probe according to the manufacturer’s recommendations (Applied Spectral Imaging) and a minimum of 10 metaphases analyzed (20,21). Additionally, a minimum of 20 DAPI-banded metaphases was analyzed to better define the chromosomal breakpoints and intrachromosomal rearrangements. Karyotyping was conducted by the MSK Cytogenetics Core Facility and all metaphases were fully karyotyped.

### Genomic characterization

Cell lines and PDX were profiled by the MSK-IMPACT (Integrated Mutation Profiling of Actionable Cancer Targets) platform, which is a large panel sequencing (NGS) assay, that was used here to detect mutations and copy-number alterations involving up to 468 cancer-associated genes (22). As the corresponding patient-matched normal DNA was unavailable, single nucleotide variants (SNVs) representing known COSMIC somatic mutations or truncating mutations in tumor suppressor genes and copy number variants (CNVs) were tabulated.

### Viability and caspase 3/7 assays

For viability assays, cells were plated in clear-bottom, white 96-well plates at a density of 5,000 cells per well and incubated with compounds for 96 h. The relative amount of viable cells was determined using AlamarBlue viability dye and fluorescence was measured using a SpectraMax M2 plate reader (Ex: 485 nm, Em: 530 nm) (23). Data was analyzed by non-linear regression and curves fitted using GraphPad Prism software to generate IC_50_ values. For caspase 3/7 activity, cells were plated at a density of 30,000 cells/well directly into inhibitors in white, clear-bottom 96-well plates, grown for 48 h and then caspase 3/7 enzymatic activity determined using Apo-One Homogenous caspase 3/7 activity assay kit (Promega) following the manufacturer’s instructions. All data is expressed relative to control values and is an average of 2-5 independent experiments where each condition was assayed in triplicate determinations.

### Plasmids, lentiviral generation and transfection

Bacteria stocks harboring the pLKO.1 MISSION lentiviral shRNA constructs were obtained from the MSK RNAi Core Facility. For packaging lentivirus, we used psPAX2 (Addgene) and VSV-G/pMD2 (Addgene). Viruses were generated and cells infected as we have described previously (24). LP9 cells were transfected with pCDNA3.1 harboring *EWSR1-WT1* fusion cDNA (cloned from the JN-DSRCT-1 cell line) or empty plasmid and 48 h later cells were treated with 500 μg/mL geneticin to select stable cells. Stable cells were obtained after 3 weeks of antibiotic selection.

### Preparation of whole-cell extracts immunoprecipitation and Western blotting

Protein levels and phosphorylation state were detected by Western blotting. For studies of ligand-dependent activation of signaling pathways, cells were deprived of serum for 24 h by growing in KSM-defined media (serum-free medium) and then treated with ligands for 15 min in KSM-defined media. Cells were lysed in 1X RIPA lysis buffer containing Halt protease and a phosphatase inhibitor cocktail according to manufacturer’s protocol (ThermoFisher Scientific). For immunoprecipitation studies, whole-cell extracts were prepared as described above and then EGFR was immunoprecipated from 300 μg total protein using an anti-EGFR antibody conjugated to agarose beads overnight. Immunoprecipitates were washed 4 times with ice-cold RIPA buffer containing protease and phosphatase inhibitors and then bound protein eluted with 100 μM 2X Laemmli sample buffer (LSB) with 5% reducing agent. Lysates or immunoprecipitates were denatured in 2X sample buffer at 55°C for 15 min, resolved on 4–12% NuPAGE gels (Invitrogen) and transferred onto PVDF (polyvinylidene fluoride) membranes. Membranes were blocked in 3% bovine serum albumin (BSA) in tris-buffered saline supplemented with 0.1% Tween-20 (vol/vol) for 1 h at room temperature and probed with primary antibodies with specificity as outlined in **Supplementary Table S1**. Proteins were separated on 8% NuPAGE gels for detection of WT1. Bound antibodies were detected with peroxidase-labeled goat antibody to mouse IgG or rabbit IgG (R&D Systems) and developed with enhanced chemiluminescence (ECL) Western blotting detection reagent (GE Healthcare).

### Proteome profiling arrays

We used human proteome profiling arrays (R&D Systems) that contain duplicate validated controls and capture antibodies that can simultaneously detect the phosphorylation state of 43 human kinases (Proteome Profiler Human Phospho-kinase Array kit) or 49 receptor tyrosine kinases (Proteome Profiler Human Phospho-RTK Array kit). Five million cells were plated in 10-cm dishes and grown for 48 h. Cells were then deprived of serum overnight and detection of protein phosphorylation was carried out according to the manufacturer’s instructions. In brief, the array membranes were blocked, incubated with 350 μg total cellular protein per array overnight at 4°C on a rocking platform, washed, and incubated with antibodies. Captured phosphorylated proteins were detected by ECL and imaged on x-ray films. The average pixel densities of duplicate spots were measured using ImageJ software (http://imagej.nih.gov/ij/), and is expressed relative to the positive control on each array.

### Detection of *EWSR1/WT1* fusion and qPCR for *WT1* and *ERBB* pathway gene expression

**Total** RNA was extracted using a Qiagen RNA mini kit and cDNAs were synthesized using SuperScript IV VILO (ThermoFisher) according to the manufacturers’ instructions. The *EWSR1/WT1* fusion was detected by RT-PCR using 5’-CTATTCCTCTACACAGCCGACT-3’ (forward, *EWSR1* exon 7) and 5’-CTGTATGTCTCCTTTGGTGTCT-3’ (reverse, *WT1* exon 8) and following conditions in a PCR thermocycler: initial denaturation (95 °C, 5 min) was followed by 38 cycles of denaturation (95 °C, 30 s), annealing (60 °C, 30 s), and extension (72 °C, 1 min); the reaction was finished after the final extension step (72 °C, 5 min). For qPCR expression analysis, Taqman assays were used and these are detailed in **Supplementary Table S1**.

### Histology and Immunohistochemistry

Histology and immunohistochemistry (IHC) were performed as previously described (25). Briefly, xenograft tissues were collected, fixed in 4% buffered formalin-saline at room temperature for 24 h, embedded in paraffin blocks and then sections of 4 μM thickness were mounted on glass slides. For IHC assays, slides were immersed in 3% H_2_O_2_ for 5 min, washed, then 15 min in 5% bovine serum albumin to block nonspecific binding sites and finally incubated in primary antibodies overnight at 4oC. The slides were washed the next day and then incubated with biotinylated anti-rabbit secondary antibody using a DAB kit (Dako) according to the manufacturer’s protocol. Slides were counterstained with hematoxylin. An anti-WT1 antibody that was raised against a peptide mapping at the C-terminal region of WT1 (C-19, Santa Cruz Biotechnology) was used.

### Establishment of patient-derived cell lines and xenografts, and efficacy studies

Patient-derived cell lines and xenografts were developed under institutional review board approved biomarker specimen protocols (06-107, 14-091) and all patients consented to collection of tumor material. All mice were cared for in accordance with guidelines approved by the Memorial Sloan Kettering Cancer Center Institutional Animal Care and Use Committee and Research Animal Resource Center and animals were monitored daily. Pleural effusion, ascites and tumor samples were collected from routine procedures in sterile containers. Heparin was added to fluids at time of collection and cells were collected by centrifugation (1000 RPM, 5 min), washed once with ice-cold PBS and then either resuspended in growth media (DMEM:F12 + 10% FBS + 1% antibiotic/antimycotic solution) or prepared as described next for solid tumors for implantation into mice. Fresh tumor samples were cleaned and then minced, mixed with Matrigel and implanted into a subcutaneous flank of female *NOD/SCID* gamma (NSG, Jackson Laboratory, Bar Harbor, ME) mice to generate xenografts (26). For cell line xenografts, 10 million cells were mixed with Matrigel (1:1) and injected subcutaneously into a single flank of female NSG mice. Once tumors reached approximated 100 mm^3^ volume mice were randomize into groups of 4 or 5 mice to achieve similar average starting tumor volume across groups and tumor-bearing animals were treated with vehicle, afatinib (25 mg/kg QD), cetuximab (1 mg BIW), afatinib (25 mg/kg QD) + cetuximab (1 mg BIW) when tumors reached approximately 100 mm^3^ volume. Afatinib was resuspended in 0.5% methylcellulose and 0.4% Tween 80) and was given by oral gavage. Cetuximab was administered via intraperitoneal injection. Tumors size and body weight were measured twice weekly and tumor volume was calculated using the formula: length x width^2^ × 0.52.

### Microarray expression analysis

We mined legacy gene expression microarray data for 137 sarcoma samples (28 DSRCT, 28 Ewing sarcomas (ES), 23 fusion-positive alveolar rhabdomyosarcomas (ARMS), 46 synovial sarcomas (SS) and 12 alveolar soft-part sarcomas (ASPS)) that were generated using Affymetrix U133A arrays and used in previous publications (9,10,27). The raw data are available at http://cbio.mskcc.org/public/sarcoma_array_data/filion2009.html. Data were Robust Multichip Average (RMA)-normalized and then linear regression analysis was performed using limma R package (28). Gene Set Variability Analysis was conducted with GSVA R package (29) and GSEA enrichment analysis was performed using java-based GSEA software (29). The GSVA computes enrichment scores for predefined gene sets (oncogenic signatures) for each sample separately, allowing for more flexible comparison of multiple groups and assessing of intragroup variability. The average scores in each group were then compared between sarcoma types using linear regression in limma (28). Exploratory analysis was performed using Rtsne and pheatmap packages. GSEA analysis was run using “gene_set” permutation type against the “Oncogenic Signatures” gene sets database. pheatmap, ggplot2 and clusterProfiler R packages were used for data visualization (30).

### Statistical Analysis

There were two to three replicates of each condition and all experiments were repeated 2-4 times. For animal studies there were 5 mice in each group and area under curve (AUC) analysis was used to compare the average tumor volume between groups. Briefly, area under curve values and their standard errors were computed as an estimation of a surface area between baseline values (mean value of the tumor volumes at the beginning of the treatment) and growth curves for vehicle and each treatment conditions. Treatment response was compared to the vehicle group using ANOCA with multiple Student’s t-tests. Student’s t-test was used to compare caspase activity or protein phosphorylation. For animal studies. All data were plotted and analyzed using GraphPad Prism software.

## Results

### Transcriptome profiling identifies the *ERBB* Pathway as activated in DSRCT tumor samples

To discover oncogenic pathways that may be selectively activated in DSRCT, we mined legacy microarray-based mRNA expression profiling data of 137 translocation sarcoma samples (27). This dataset comprised 28 DSRCT, 28 Ewing sarcomas (ES), 23 fusion-positive alveolar rhabdomyosarcomas (ARMS), 46 synovial sarcomas (SS) and 12 alveolar soft-part sarcomas (ASPS), all fusion-verified by clinical RT-PCR assays. These sarcomas are characterized by distinct chromosomal translocations which produce chimeric transcription factors whose unique composition results in specific signatures of pathway activation. An unbiased clustering of the expression data using t-SNE analysis (31) organized samples into clearly defined groups according to sarcoma type (**Figure 1A**), confirming the highly distinctive transcriptomes generated by these aberrant transcription factors.

**Figure 1.**
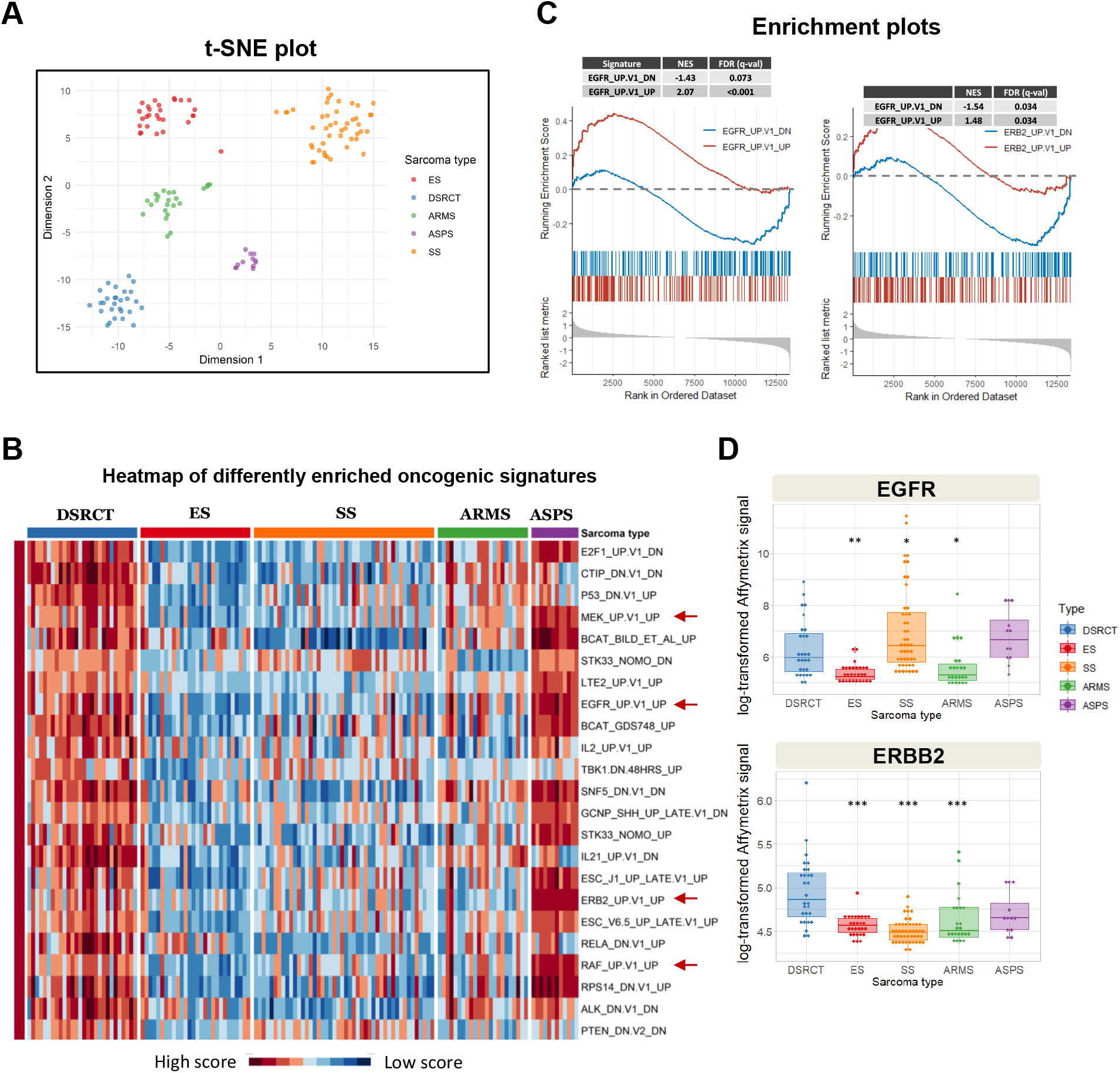
Analysis of mRNA expression data in 137 sarcoma samples reveals activation of the ERBB pathway in DSRCT. (**A**) t-SNE analysis. Sarcoma type-specific clustering of samples supports intergroup differential expression analysis. (**B**) GSVA analysis heatmap. GSVA is a non-parametric unsupervised method that allows the assessment of gene set enrichment in each individual sample. “Oncogenic signatures” from the Molecular Signature Database were queried. Enrichment scores were then compared in sarcoma types using linear regression and the top gene sets differentially enriched in DSRCT relative to other sarcomas are presented as a heatmap. (**C**) GSEA analysis was also performed comparing DSRCT to ES samples using “Oncogenic signatures”. Enrichment plots for *ERBB*-related pathways are presented with normalized enrichment scores (NES) and False Discovery Ratio (FDR, q-value). (**E**) Box plots of the expression of *EGFR* and *ERBB2*. Log_2_-transformed raw values are presented. ***Adjusted p value < 0.0001, **adjusted p value = 0.008, *Adjusted p value = 0.08.

To determine if DSRCT has a distinct gene expression profile reflecting activated growthpromoting pathways that could be exploited for therapy, we interrogated the transcriptomes of the samples from the 5 sarcoma types. We utilized the Gene Sets Variability Analysis (GSVA) with oncogenic signature gene sets from Broad MsigDB to identify activated oncogenic pathways. The top oncogenic signatures that were significantly up- or down-regulated in DSRCT compared to each of the other sarcomas were ranked by adjusted p-value and is depicted in **Figure 1B**. The finding of EGFR_UP.V1_UP and ERBB2_UP.V1_UP gene set enrichment in DSRCT compared to other sarcomas suggested a role for EGFR and ERBB2 pathway activation in these tumors. In addition, there was significant upregulation of *RAF* and *MEK* signatures, two principal ERBB family effector pathways. These results were also apparent with gene set enrichment analysis (GSEA) of DSRCT compared to ES samples using Oncogenic Signatures from Broad MsigDB which also showed that the *EGFR* pathway is active in DSRCT tumors and underscore the importance of *ERBB* pathways in DSRCT (**Figure 1C**).

Given that EGFR functions either as a homodimer or as a heterodimer with other members of the ERBB family of receptors (32), we compared expression of the four *ERBB* family genes in DSRCT relative to the other four sarcomas in the dataset. Among the five sarcomas, only DSRCT showed relatively high mRNA expression of both *EGFR* and *ERBB2* (**Figure 1D**)., supporting a potential role for the ERBB family receptors in DSRCT growth. DSRCT tumors did not have a significantly higher level of expression of *ERBB3* or *ERBB4* mRNA compared to the other sarcomas (data not shown).

### Generation of novel tumorigenic DSRCT cell lines

Investigation into the biology of DSRCT has been hindered by a paucity of preclinical disease models. Only the JN-DSRCT-1 cell line is well characterized and widely used (16). To address this obstacle, we sought to develop new patient-derived DSRCT cell lines and xenograft models. We generated two DSRCT cell lines from Memorial Sloan Kettering patient samples (SK-DSRCT1, SK-DSRCT2) and characterized two DSRCT cell lines that were previously published but not genomically or biochemically characterized (BER-DSRCT, BOD-DSRCT) (18). The morphology of the cells is shown in **Figure 2A**, with images obtained at two different degrees of confluence and patient demographic information is provided in **Supplementary Table S2**. The BOD-DSRCT and BER-DSRCT cells appear as small round cells at low and high densities, similar to JN-DSRCT-1 cells. SK-DSRCT1 and SK-DSRCT2 cells appear as mixed morphology with both cell lines having mixed round and spindle-shaped cells.

**Figure 2:**
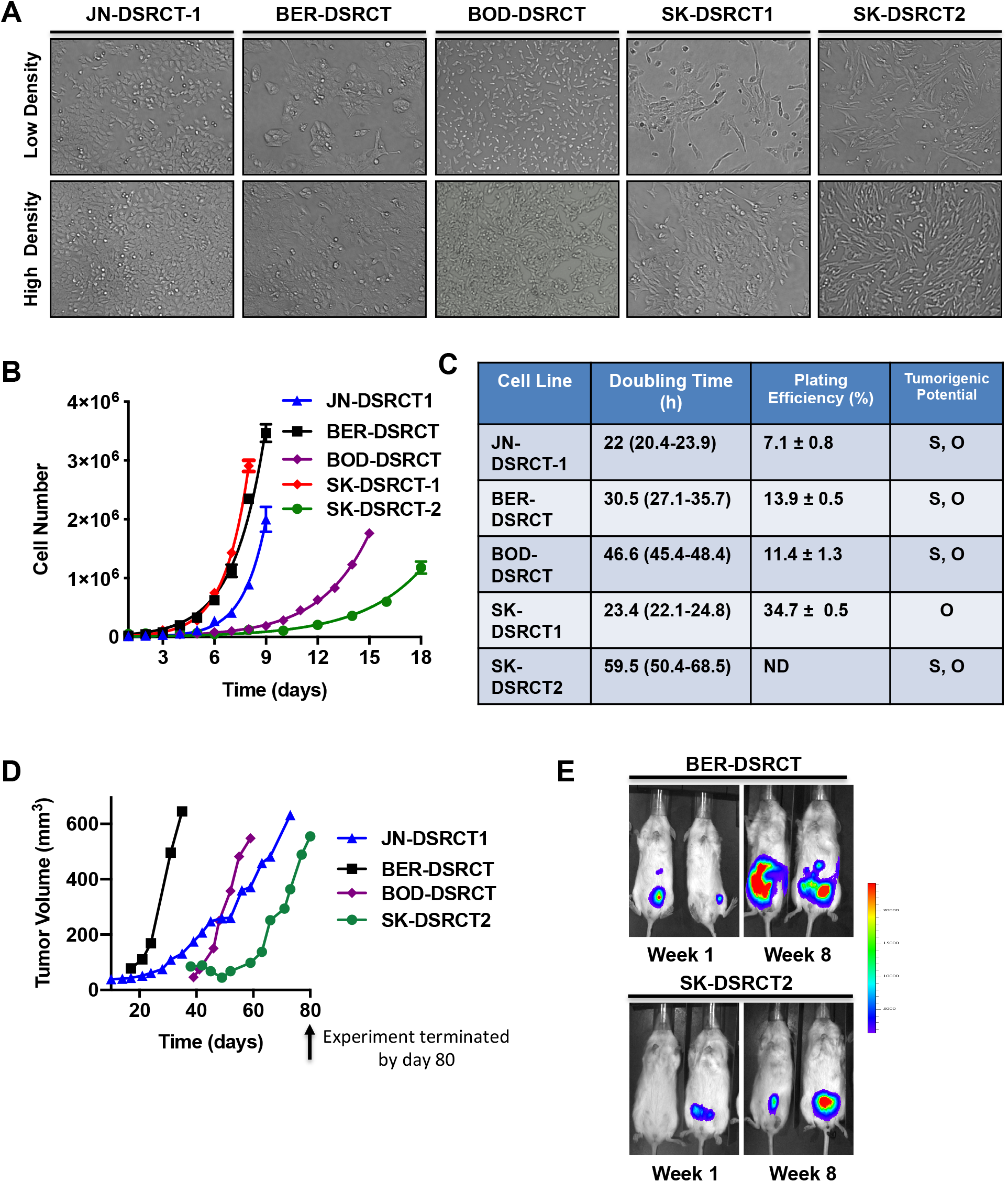
Generation and characterization of novel DSRCT preclinical models. (**A**) Phase-contrast images of DSRCT cell lines (100X magnification). (**B**) Growth characteristics of DSRCT cell lines in culture. (**C**) Doubling time, plating efficiency and tumorigenic potential of cells are shown in the right panel. (**D**) DSRCT cell lines were implanted into the subcutaneous flank of immunocompromised mice and tumors were measured twice weekly. Data represent the average volume of two tumors per cell line. (**E**) Cell lines stably expressing a luciferase construct were implanted into the peritoneal cavity of immunocompromised mice, and bioluminescence images were acquired weekly. The first (one week after implantation) and last (8 weeks after implantation) images are shown. S: subcutaneous. O: orthotopic.

We next characterized the growth rate, plating efficiency and tumorigenic potential of the five DSRCT cell lines. The growth rate of cell lines was variable with JN-DSRCT-1 and SK-DSRCT1 having the shortest doubling time (22 and 23.4 h, respectively), followed by BER-DSRCT (**Figure 2B and 2C**). The SK-DSRCT2 and BOD-DSRCT cell lines were slowest growing with doubling times of 59.5 and 46.6 h, respectively (**Figure 2B and 2C**). In clonogenic assays, the JN-DSRCT-1 cell lines had the lowest plating efficiency (7.1 ± 0.8%) and the SK-DSRCT1 cell line had the highest plating efficiency (34.7 ± 0.5%) (**Figure 2C**).

To explore the ability of the cell lines to form xenograft tumors, cells were implanted either in the subcutaneous flank (unlabeled cells) or the peritoneal cavity (cell lines stably expressing a GFP-luciferase cDNA) of immunocompromised mice. Following subcutaneous implantation, all cell lines except SK-DSRCT1 grew into xenograft tumors by day 80 (when the study was terminated) (**Figure 2D**). For the orthotopic xenograft models, luciferase-expressing BER-DSRCT and SK-DSRCT1 cells were injected into the peritoneum and then bioluminescence images were acquired weekly. Images taken one and eight weeks after implantation of cells in two mice per cell line are shown in **Figure 2E**. The luciferase signals increased over this time, indicating that both BER-DSRCT and SK-DSRCT1 (which did not form subcutaneous tumors) are capable of forming orthotopic xenograft tumors in the peritoneal cavity, which is the most common site of presentation of DSRCT in patients.

### Cytogenetic and genomic characterization of novel DSRCT cell lines

Cytogenetic analysis was performed on all cell lines to examine the karyotype. The chromosome spread obtained by either SKY karyotyping or DAPI banding are shown in **Figure 3A** (SK-DSRCT1 cell line) and in **Supplementary Figure S1** (all cell lines). More details of the cytogenetic analysis of each cell line are presented in **Supplementary Table S3**. The t(11;22)(q13;q12) was identified in all five cell lines, with the typical balanced translocation in four cell lines. The SK-DSRCT1 cell line was the only one with an unbalanced t(11;22)(q13;q12). Four of the five DSRCT cell lines had gained either an extra chromosome 5, or had duplication of some region or some parts of it (only BOD-DSRCT showed no copy number alteration of chromosome 5). There were also gains and losses and additional rearrangements that were unique to each cell line (**Supplementary Table S3**).

**Figure 3:**
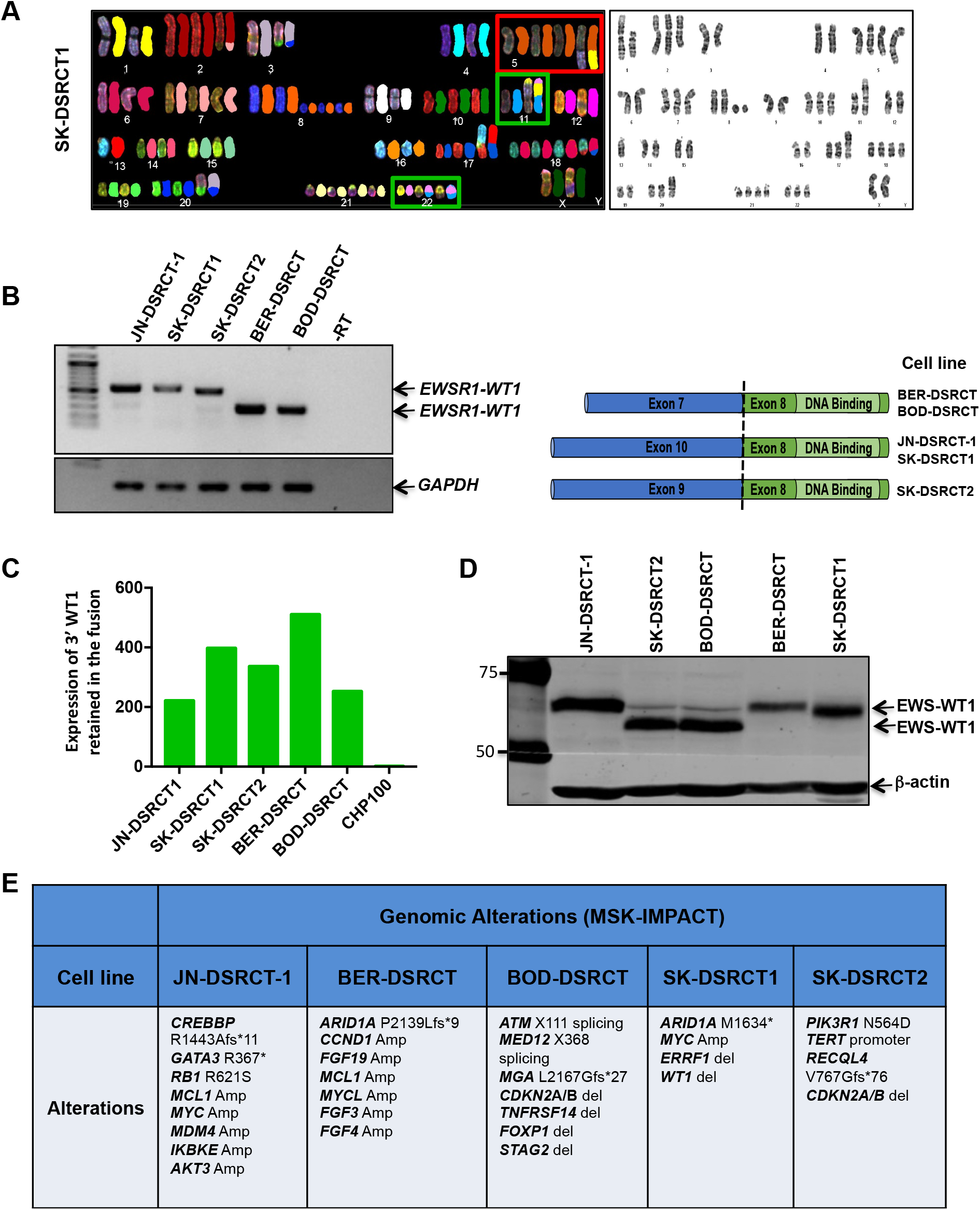
Cytogenetic, genetic and genomic characteristics of novel DSRCT cell lines. (**A**) Spectral (left) or DAPI-banded (right) karyotypes of SK-DSRCT1 cells illustrating the translocation between chromosomes 11 and 22, resulting in the oncogenic chimeric transcription factor, *EWSR1-WT1* is shown in the green box. Chr5 polysomy is shown in the red box. The karyotypes of all cell lines are shown in Supplementary Figure 1. (**B**) RT-PCR conducted on five DSRCT cell lines showing expression of the *EWSR1-WT1* fusion (left). PCR amplicons were TOPO-cloned and then validated by orthogonal DNA sequencing. The exons of *EWSR1* and *WT1* that were fused in each cell line are shown in the cartoon (right). (**C**) Expression of 3’ *WT1* mRNA retained in the fusion. *EWSR1-WT1* mRNA level is expressed relative to that of native *WT1* in CHP100 Ewing sarcoma cell line. (**D**) Western blot analysis of EWS-WT1 using an anti-WT1 antibody targeting the C-terminal region of WT1 in DSRCT cell lines. (**E**) DNA was profiled by the MSK-IMPACT platform to identify genomic alterations in cell lines.

To confirm expression of the chimeric *EWSR1-WT1* gene resulting from the t(11;22)(q13;q12) translocation, we performed RT-PCR using primers for *EWSR1* exon 7 and *WT1* exon 8. Three cell lines had amplicons of similar sizes (JN-DSRCT-1, SK-DSRCT1, SK-DSRCT2) and the remaining two cell lines had a smaller sized product (**Figure 3B, left panel**). PCR amplicons were cloned into a TOPO vector, expanded in bacteria and then plasmid DNA from ten colonies was isolated and the inserts were sequenced to identify the precise exons of *EWSR1* and *WT1* that are fused in each cell line. A cartoon of the results are shown in **Figure 3B, right panel**. The cell lines harbored heterogeneous in-frame fusions between *EWSR1* (various exons) and exon 8 of *WT1* (**Figure 3B, right panel**), as has been shown for DSRCT tumors (33,34). The BER-DSRCT and BOD-DSRCT cell lines had a fusion between *EWSR1* exon 7 and *WT1* exon 8, which as we have shown previously, is present in the majority of DSRCT tumors (4). The other cell lines, including JN-DSRCT-1, had *EWSR1-WT1* fusions that occur less frequently in patient samples. Although a “9-8” fusion transcript was predominant in the SK-DSRCT2 cell line, it also expressed a very low level of the “7-8” transcript, presumably from the same allele (**Figure 3B**).

We further validated *WT1* expression by quantitative PCR (qPCR) targeting the 3’ part of *WT1* that is retained in the fusion and Western blotting using an anti-WT1 antibody that was raised against a peptide mapping at the C-terminal region. The Ewing sarcoma cell line CHP100 (EWS-FLI1 fusion) as a negative control. As expected, *WT1* mRNA was detected in all DSRCT cell lines and amounted to 200-500-fold higher than in CHP100 (**Figure 3C**). Western blotting with the C-terminal WT1 antisera confirmed the presence of WT1 in all of the new DSRCT cell lines (**Figure 3D**). The band detected is most likely EWS-WT1 as transcriptomic analysis has shown that the JN-DSRCT-1 cell line does not express wildtype WT1 (35).

To broadly characterize our novel cell line models, we profiled DNA by NGS using our MSK-IMPACT (<u>i</u>ntegrated <u>m</u>utation <u>p</u>rofiling of <u>a</u>ctionable <u>c</u>ancer <u>t</u>argets) platform which captures genomic alterations across 468 genes of proven or potential therapeutic and/or prognostic significance (22). Known COSMIC somatic mutations or truncating mutations in tumor suppressor genes as well as CNVs are listed for each DSRCT cell line in **Figure 3E**. Among the notable findings was a *STAG2* deletion in one sample, a gene known to be recurrently inactivated in an aggressive subset of Ewing sarcomas (36), another *EWSR1*-rearranged sarcoma (37).

### Proteomic profiling of DSRCT models identifies elevated ERBB family RTK activity

The results presented in **Figure 1** indicate that transcriptional signatures of activated kinase signaling pathways are present in DSRCT. To determine which of these pathways are activated at the proteomic level, we profiled the phosphorylation state of 49 RTKs using phospho-proteomic arrays in the five DSRCT cell lines and five DSRCT patient tumors (**Figure 4A-D**). Protein phosphorylation was quantitated by densitometry and is shown in **Figure 4B** and **D**. We found that members of the ERBB family RTKs were phosphorylated in all cell lines, with phosphorylation of EGFR, ERBB4 and ERBB2 consistently high in all cell lines (**Figure 4A**). Note that due to the use of different primary antibodies on the array, the level of absolute level of phosphorylation of each ERBB RTK cannot be compared to each other. Only phosphorylation of EGFR was consistently observed in multiple DSRCT cell lines and tumors (**Figure 4B and 4D**). To validate these data, we performed immunoblotting using phospho-specific anti-ERBB family antibodies, using the untransformed mesothelial cell line LP9 (17) as a control. Phosphorylated EGFR, ERBB2, and ERBB4 were detected in all cell lines, and consistent with the low signal in the RTK proteomic arrays, phosphorylated ERBB3 was extremely low (**Figure 4E**). To determine if these activated ERBB RTKs interacted with each other, we immunoprecipitated EGFR and then immunoblotted for each ERBB RTK using JN-DSRCT-1 cell extracts (**Figure 4F**). We were able to clearly detect ERBB2/HER2 and ERBB4/HER4 in anti-EGFR immunoprecipitates, suggesting that these three RTKs interacted or co-localized. There was also a very weak band on the ERBB3 Western blot, indicating that there may also be some interaction between EGFR and ERBB3.

**Figure 4:**
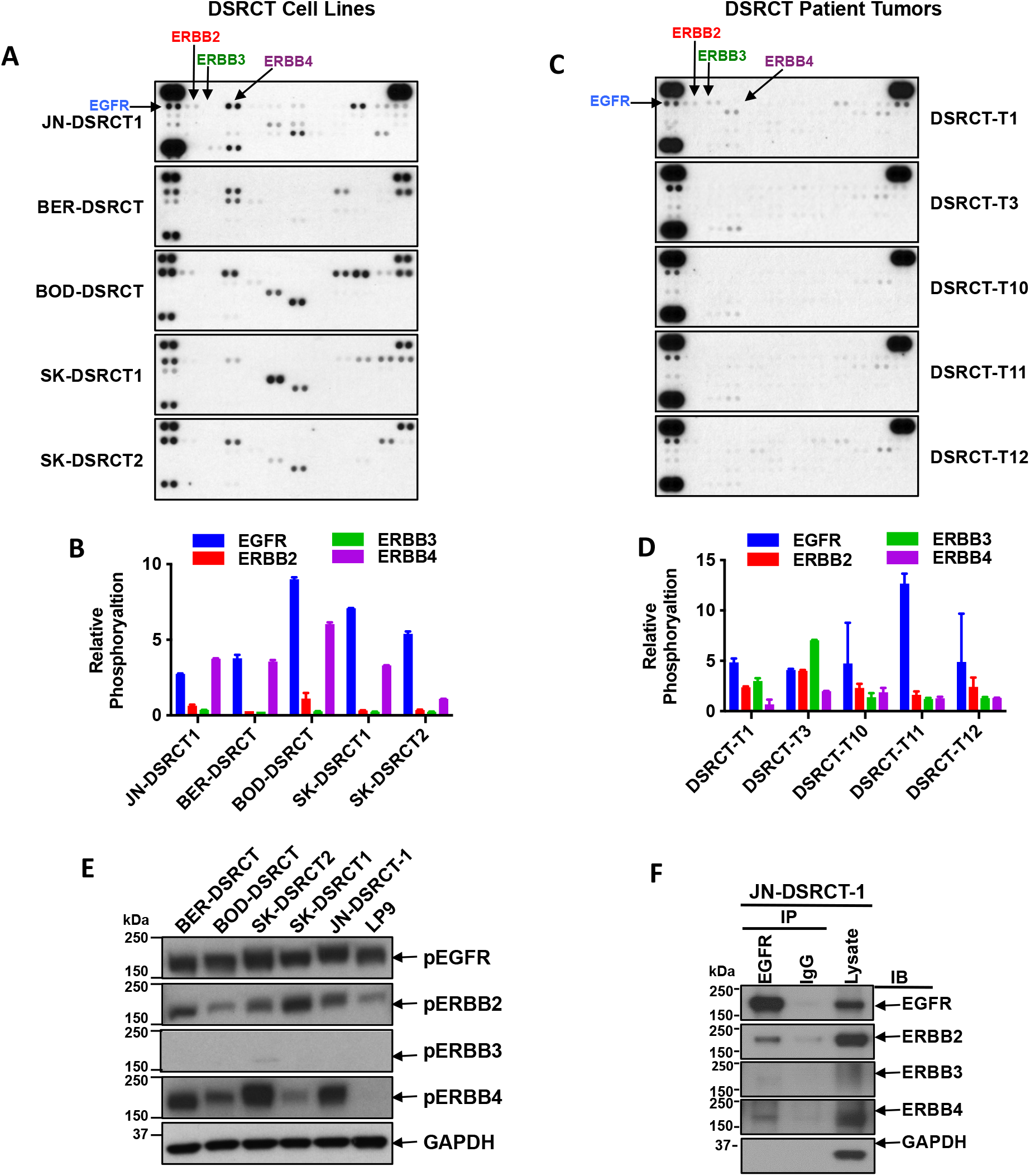
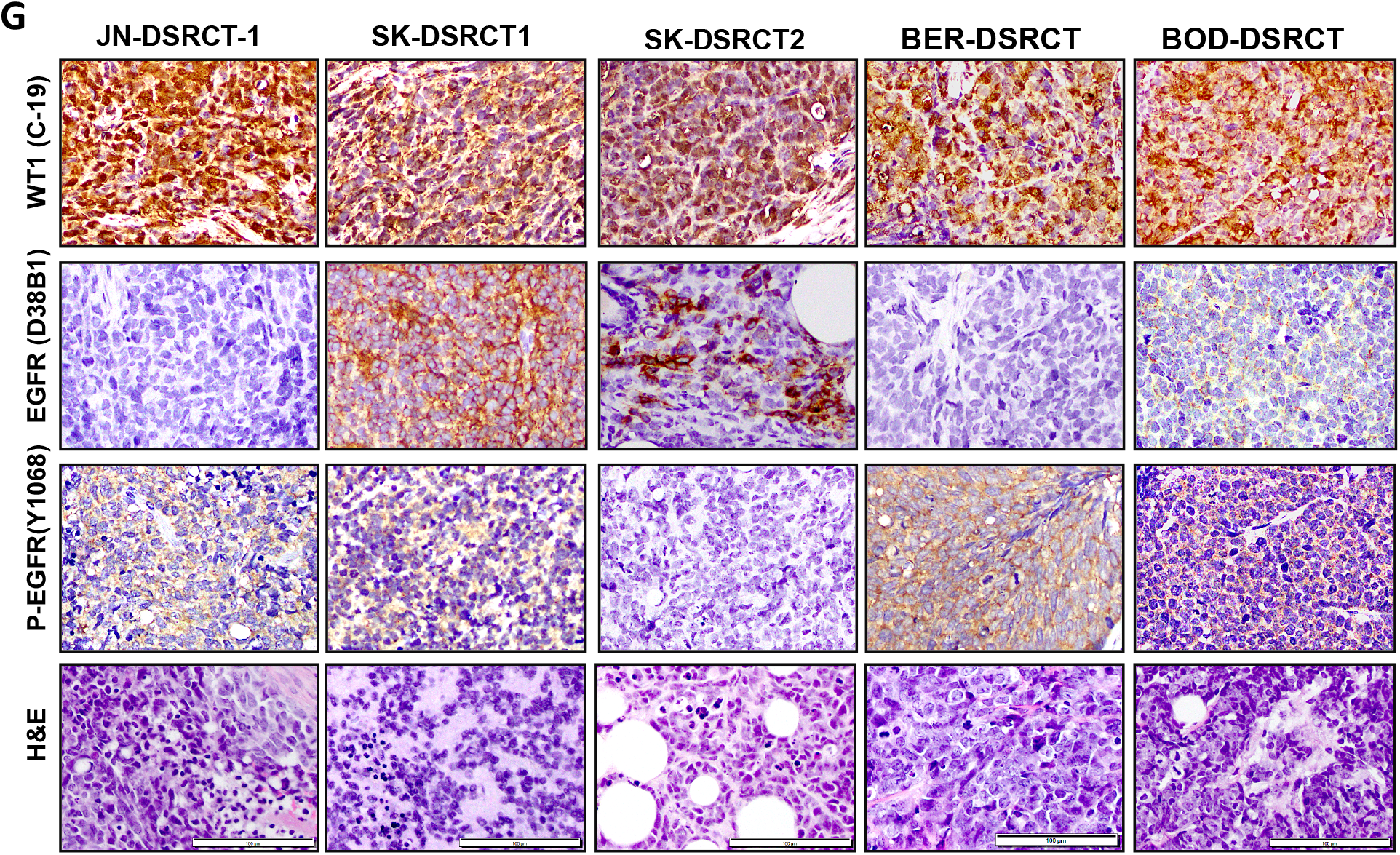
Expression and activation of ERBB family members in novel DSRCT cell lines. Receptor tyrosine kinase (RTK) arrays were used to profile phosphorylated RTKs in DSRCT cell lines (**A and B**) or tumors (**C and D**). Proteomic arrays were imaged on x-ray films and quantitated by densitometry. Values are expressed relative to values obtained in CHP100 cells (ES cell line). (**E**) Phosphorylation of ERBBs was examined by Western blot using phosphospecific antibodies. LP9: non-tumorigenic mesothelial cell line. (**F**) EGFR was immunoprecipitated from JN-DSRCT-1 cells and then Western blot analysis conducted to identify other ERBB family RTKs that are associated with it. (**G**) Detection of WT1, EGFR and phospho-EGFR by immunohistochemistry in paraffin-embedded xenograft tumors or cell pellet (SK-DSRCT1). Representative photomicrographs are shown. Scale bar: 100 μm.

We next performed immunohistochemistry on paraffin-embedded DSRCT cell line xenografts or cell pellets (SK-DSRCT1) to confirm the presence of WT1 using an antibody raised to the C-terminus of WT1, and to explore expression of EGFR and phosphorylated EGFR, given that this ERBB RTK was the most consistently activated in cell lines and tumors tested. Strong nuclear staining for the WT1 C-terminus was detected in all samples probed, most likely representing EWSR1-WT1 in these novel DSRCT models, as a recent transcriptome profiling study of has confirmed a lack of native *WT1* transcripts in DSRCT (35). EGFR was detected at varying levels in the different DSRCT patient xenografts or xenografted DSRCT cell lines. The EGFR staining of JN-DSRCT1 xenograft tissue is consistent with the EGFR IHC staining that we have previously obtained in a JN-DSRCT-1 cell block (38). However, phosphorylated EGFR was detected in all samples, consistent with our *in vitro* studies (**Figure 4G**).

### The ERBB pathway regulates growth of DSRCT cells

To explore the relationship between expression of EWS-WT1 and the ERBB pathway in an isogenic system, we looked at the level of components of the *ERBB* pathway following expression of EWS-WT1. A cDNA encoding *EWSR1-WT1* was stably transfected into LP9 cells and then expression of the mRNAs for *ERBB* RTKs and ligands was determined in LP9-empty vector and LP9-EWS-WT1 cells using a custom qPCR array (**Supplementary Table S2**). Expression of *EWS-WT1* in LP9 cells is shown in **Figure 5A**. EWS-WT1 expression in LP9 cells caused a significant increase in mRNAs for several ERBB pathway ligand genes, including *NRG1, EGF, AREG* and *EPGN* (**Figure 5B**). There was no change in *EGFR* mRNA, whereas *ERBB2* mRNA increased after EWS-WT1 expression and there was a significant decrease in *ERBB3* mRNA (**Figure 5B**). These results led us to hypothesize that ligands of the ERBB pathway can drive growth of DSRCT cells.

**Figure 5:**
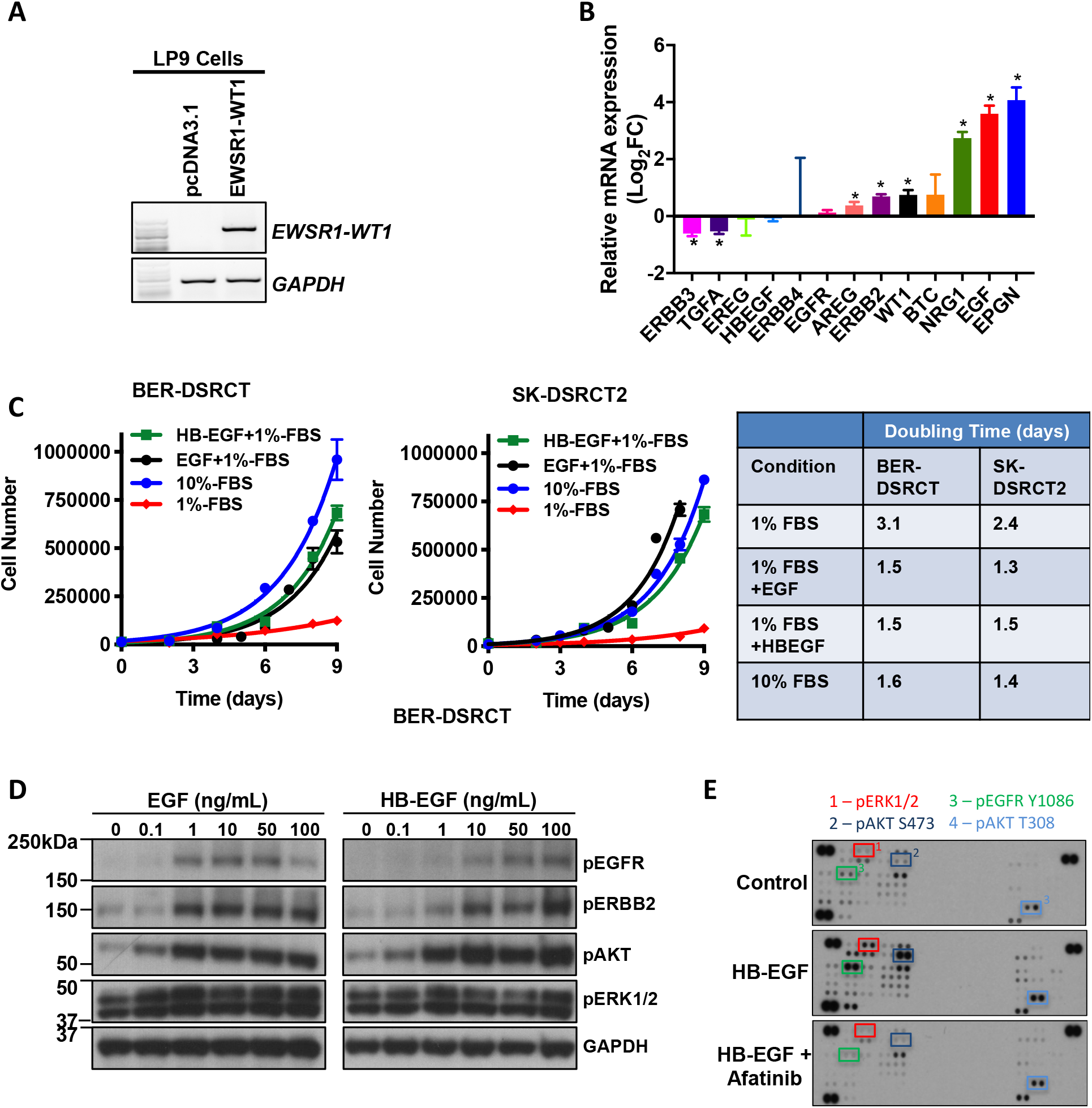
ERBB pathway genes are altered by expression of EWSR1-WT1 and regulates growth of DSRCT cell lines. A cDNA encoding *EWSR1-WT1* was expressed in LP9 cells (**A**) and then expression of known *ERBB* receptors and ligands were determined by qPCR (**B**). Expression of *EWSR1-WT1* was confirmed by RT-PCR. mRNA levels in LP9-EWS-WT1 cells are expressed relative to that in LP9-pcDNA3.1 (empty plasmid). * P < 0.05. (**C**) EGFR ligands are sufficient to stimulate growth of DSRCT cell lines. Results represent the mean ± SD of two independent experiments. Growth data was fitted to an exponential growth equation using GraphPad Prism 7 software and the doubling time is given in the right panel. (**D**) BOD-DSRCT cells were serum-starved for 24 h and then stimulated with the indicated concentrations of EGF or HB-EGF for 15 min. Western blotting was then performed for the phosphorylated proteins shown or GAPDH. (**E and F**) BER-DSRCT cells were pretreated with DMSO or 0.25 μM afatinib for 30 min and then stimulated with 100 ng/mL HB-EGF for 15 min. Phosphokinase arrays were then used to assess the phosphorylation state of selected signaling proteins. (**E**) Representative images of phosphokinase arrays. (**F**) Arrays were quantitated by densitometry and the relative change in phosphorylation above DMSO-treated control cells are shown in Supplementary Figure 2.

To address this, we treated two DSRCT cell lines with EGF or HB-EGF and examined growth and signaling pathways for activation. Chronic treatment of cells with EGF or HB-EGF was sufficient to drive a significant increase in cell growth, achieving the same level as seen in cells grown in 10% FBS (**Figure 5C**). Treatment of DSRCT cell lines with EGF or HB-EGF stimulated phosphorylation of EGFR and ERBB2 in a dose-dependent manner (**Figure 5D**). Concomitantly, the downstream effectors AKT and ERK1/2 were activated, as expected (**Figure 5D**).

For a more comprehensive view of activated pathways in DSRCT cells, we employed a phospho-kinase array that contains antibodies that recognize 43 cytoplasmic phospho-proteins known to be activated by growth factors, to profile the signaling pathways activated by treatment with an ERBB ligand in the absence or presence of the pan-ERBB inhibitor afatinib (**Figure 5E**). The relative protein phosphorylation was quantitated by densitometry and is presented in **Supplementary Figure 2**. Stimulation of BER-DSRCT cells with HB-EGF resulted in activation of multiple effector kinases, and this effect was mitigated by afatinib treatment.

To validate EGFR as a target for therapy in DSRCT, we examined the effect of pharmacological and genetic antagonists of EGFR on growth and survival. For these studies, we used afatinib and neratinib, two pan-ERBB inhibitors that have been shown to be active *in vitro* and *in vivo*. Treatment of BER-DSRCT (**Figure 6A, left panel**) and JN-DSRCT-1 (**Figure 6A, right panel**) cells with afatinib or neratinib inhibited growth of both cell lines in a dose-dependent manner, with half inhibitory concentration (IC_50_) values below 1 μM. This inhibition of growth was accompanied by an increase in apoptosis as treatment of cells with afatinib stimulated caspase 3/7 activity in the three DSRCT cell lines tested (**Figure 6B, left panel**). Consistent with these findings, genetic inhibition of *EGFR* using two unique lentiviral delivered short hairpin RNAs (shRNA) in BER-DSRCT cells resulted in significant reduction in cell number relative to a scrambled shRNA control (**Figure 6B, middle panel**). The reduction in cell number coincided with a large increase in apoptosis relative to control shRNA as measured by caspase 3/7 activity (**Figure 6B, right panel**). Treatment of SK-DSRCT1 and SK-DSRCT2 for 48 h with afatinib, cetuximab or a combination of both also resulted in significantly reduced cell number (**Figure 6C**). The consistency of these results confirms the sensitivity of these tumors to EGFR inhibition.

**Figure 6:**
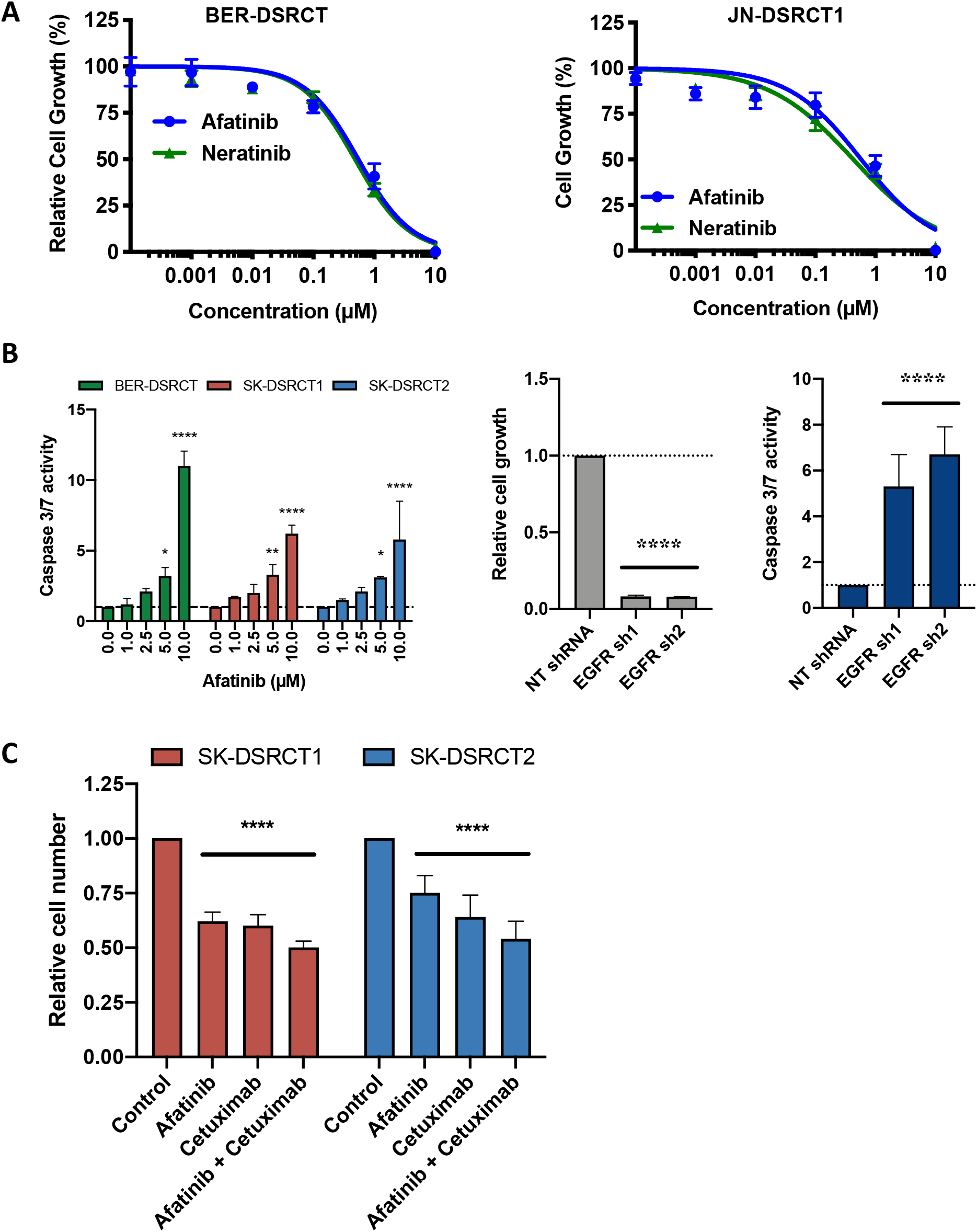
EGFR antagonists inhibit growth of novel DSRCT cell lines. (**A**) Cells were treated with increasing doses of afatinib or neratinib and then either viability or caspase 3/7 activity determined. Results represent the mean ± SD of 2-3 experiments in which each condition was assayed in 2-3 replicates. (**B**) BER-DSRCT cells were infected with lentivirus harboring shRNAs (scrambled control or targeting *EGFR*) and then the number of cells or caspase 3/7 activity determined (left panel). (**C**) Cells were treated with 1 μM afatinib,100 ng/mL cetuximab, or 1 μM afatinib + 100 ng/mL cetuximab for 48 h.****P < 0.0001.

To begin to translate these findings to DSRCT patients, we examined the efficacy of targeting EGFR in xenograft models. Cells (BER-DSRCT, SK-DSRCT2) or PDX (DSRCT-10Cpdx) tissues were implanted into the subcutaneous flank of immunocompromised mice and animals were treated with vehicle (QD), afatinib (25 mg/kg QD), cetuximab (1 mg QW), or a combination of cetuximab (1 mg QW) and afatinib (25 mg/kg QD). The tumor volumes over the course of treatment are shown in **Figure 7** (SK-DSRCT2, DSRCT-10c) and **Supplementary Figure 3A** (BER-DSRCT). We used an area under curve analysis to compare the effect of treatment between groups as this analysis takes into account the magnitude and duration of the treatment effect. Cetuximab treatment reduced growth of SK-DSRTCT2 (**Figure 7A**) and BER-DSRCT (**Supplementary Figure 3A**) tumors significantly. However, afatinib monotherapy did not reduce growth significantly of any of the cell line xenograft tumors (**Figure 7A and Supplementary Figure 3A**), even though there was a modest reduction in growth of BER-DSRCT xenograft tumors (Supplementary **Figure 3**). We extended these studies to include a DSRCT PDX model that we developed (DSRCT-10Cpdx). We confirmed that DSRCT-10Cpdx expresses the *EWSR1-WT1* fusion (**Figure 7B**). Similar to the cell lines and patient samples shown in **Figure 4**, the patient sample from which the DSRCT-10Cpdx model was derived had elevated levels of phosphorylated EGFR (**Figure 7C**). The combination of afatinib and cetuximab significantly reduced growth of this PDX model (**Figure 7D**).

**Figure 7:**
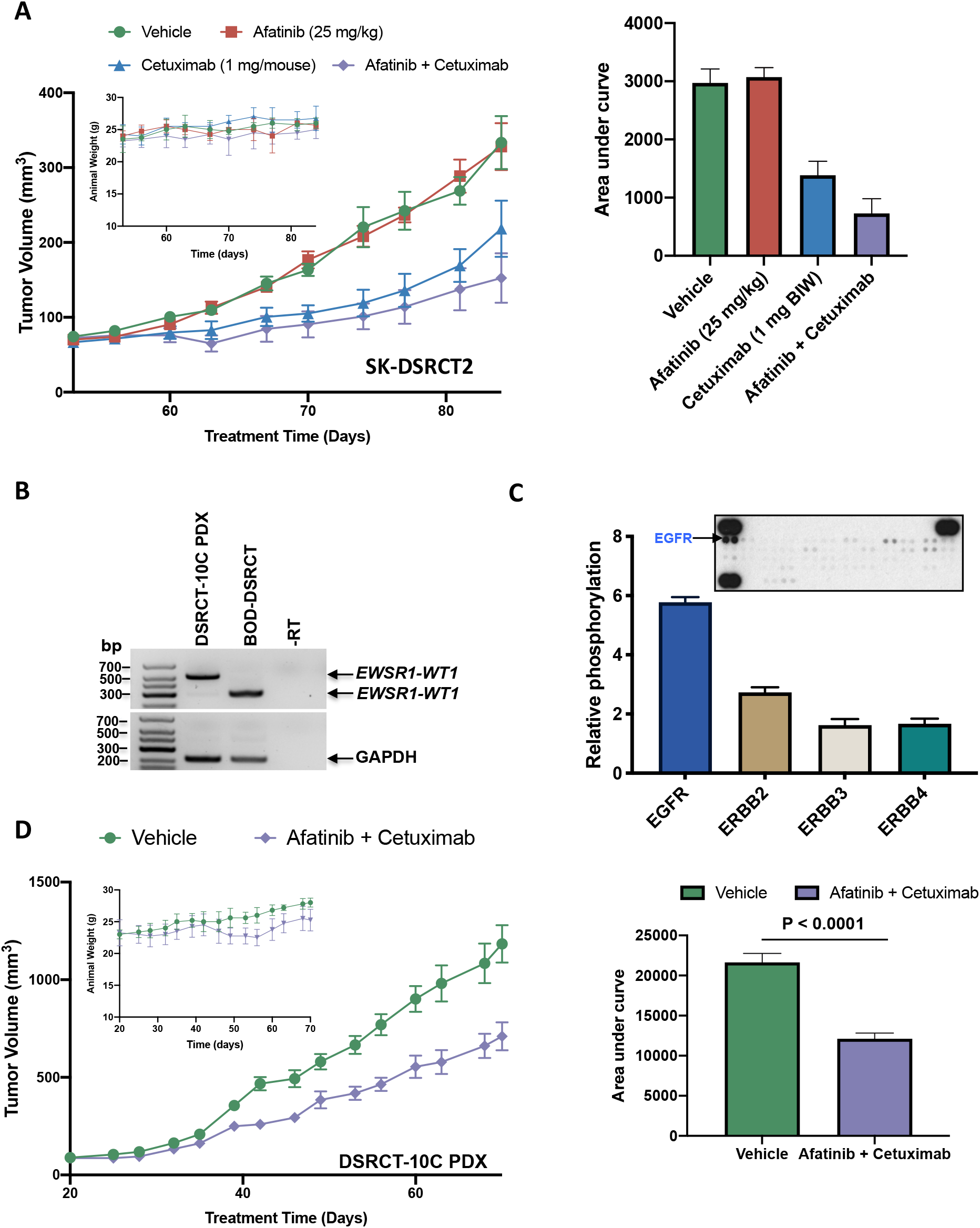
EGFR antagonists inhibit growth of novel DSRCT xenograft tumors. SK-DSRCT2 cells (**A**) or DSRCT-10C PDX tumors (**B-D**) were implanted subcutaneously into the flank of immunocompromised mice and treatment began when tumors reached approximately 100 mm^3^. Mice were treated with vehicle, afatinib (25 mg/kg, QD, 5 days/week), cetuximab (1 mg BIW), or a combination of cetuximab and afatinib. (A) Tumor volume measurements of SK-DSRCT2 xenografts with animal weight shown in the inset. Area under curve analysis is shown on the right. Expression of *EWRS1-WT1* fusion was confirmed in DSRCT-10C PDX tumor by RT-PCR (**B**). The phosphorylation level of RTKs was examined using the RTK array (**C**). The array was quantitated by densitometry and relative levels of each phosphorylated ERBB family member is shown in the accompanying graph. (**D**) DSRCT-10C PDX tumor volume is shown, demonstrating efficacy of combination therapy with afatinib and cetuximab with animal weight shown in the inset. Area under curve analysis is shown on the right.

## Discussion

The novel DSRCT models that we describe here represent an invaluable tool to the research community to investigate this aggressive sarcoma of adolescents and young adults. By analyzing transcriptomic and proteomic profiling of DSRCT tumors and cell lines, we find ERBB signaling pathways to be activated in DSRCT relative to other sarcomas. Further, we show that EGFR is a determinant of growth in these tumors and can be exploited for therapy.

Our study was focused primarily on developing new DSRCT preclinical models and to examine kinases as potential therapeutic target. Determining precisely how the ERBB and other kinase signaling pathways are dysregulated and co-opted by the EWSR1-WT1 chimeric transcription factor to drive tumor progression remains an open question and one that we can begin to investigate more effectively with these new DSRCT models.

Our previous work and that of other groups has shown that expression of *EWSR1-WT1* can activate kinase signaling, including activation of PDGF and IGF signaling (39,40). However, given the lack of DSRCT models, it was not possible to perform preclinical studies to evaluate the therapeutic relevance of those observations. Using novel DSRCT cell lines, we have approached the challenge of developing targeted therapy for DSRCT in an unbiased approach, asking which known targetable growth pathways might be dysregulated and contributing to DSRCT proliferation. We found that EGFR is consistently phosphorylated in DSRCT cell lines, PDX and patient tumors. Moreover, expression of the EWSR1-WT1 oncoprotein in LP9 cells dysregulated the ERBB pathway, increasing the expression of several ligands, including EGF, amphiregulin, epiregulin, and neuregulin 1. It is likely that the increased level of these ligands is responsible for the increased activation of EGFR (and other ERBB receptors) observed, as we also demonstrate that treating DSRCT cells with EGF is sufficient to increase proliferation.

To begin to evaluate the potential clinical relevance of our findings, we explored the effect of inhibiting EGFR in cell lines and xenograft models. Both genetic and chemical perturbation of EGFR signaling resulted in reduced growth and increased apoptosis. Extending these studies to animal models, we found that cetuximab was the most effective anti-EGFR agent for reducing tumor burden in mice bearing DSRCT tumors. Although the reduction in tumor growth obtained when EGFR antagonists were administered *in vivo* results were only partial, we believe that these finds are promising and should be further explored to determine if different dosing or combination with chemotherapy or other targeted agents would offer additional benefits for patients. Given the poor response and limited overall survival of patients with DSRCT, our findings begin to provide new therapeutic options where they are desperately needed.

## Acknowledgements

This work was partially supported by a U54 OD020355 grant from the National Institute of Health to Elisa de Stanchina and a Memorial Sloan Kettering Cancer Center Support Grant (P30 CA008748). Funding for this project was also provided by Elevation Oncology.

## References

1. Gerald WL, Miller HK, Battifora H, Miettinen M, Silva EG, Rosai J. Intra-abdominal desmoplastic small round-cell tumor. Report of 19 cases of a distinctive type of high-grade polyphenotypic malignancy affecting young individuals. Am J Surg Pathol 1991;15(6):499–513.

2. Sawyer JR, Tryka AF, Lewis JM. A novel reciprocal chromosome translocation t(11;22)(p13;q12) in an intraabdominal desmoplastic small round-cell tumor. Am J Surg Pathol 1992;16(4):411–6 doi 10.1097/00000478-199204000-00010.

3. Shen WP, Towne B, Zadeh TM. Cytogenetic abnormalities in an intraabdominal desmoplastic small cell tumor. Cancer Genet Cytogenet 1992;64(2):189–91 doi 10.1016/0165-4608(92)90355-c.

4. Gerald WL, Rosai J, Ladanyi M. Characterization of the genomic breakpoint and chimeric transcripts in the EWS-WT1 gene fusion of desmoplastic small round cell tumor. Proc Natl Acad Sci U S A 1995;92(4):1028–32 doi 10.1073/pnas.92.4.1028.

5. Gerald WL, Haber DA. The EWS-WT1 gene fusion in desmoplastic small round cell tumor. Semin Cancer Biol 2005;15(3):197–205 doi 10.1016/j.semcancer.2005.01.005.

6. Ladanyi M, Gerald W. Fusion of the EWS and WT1 genes in the desmoplastic small round cell tumor. Cancer Res 1994;54(11):2837–40.

7. Hayes-Jordan A, LaQuaglia MP, Modak S. Management of desmoplastic small round cell tumor. Semin Pediatr Surg 2016;25(5):299–304 doi 10.1053/j.sempedsurg.2016.09.005.

8. Lal DR, Su WT, Wolden SL, Loh KC, Modak S, La Quaglia MP. Results of multimodal treatment for desmoplastic small round cell tumors. J Pediatr Surg 2005;40(1):251–5 doi 10.1016/j.jpedsurg.2004.09.046.

9. Kobos R, Nagai M, Tsuda M, Merl MY, Saito T, Lae M, et al. Combining integrated genomics and functional genomics to dissect the biology of a cancer-associated, aberrant transcription factor, the ASPSCR1-TFE3 fusion oncoprotein. J Pathol 2013;229(5):743–54 doi 10.1002/path.4158.

10. Tsuda M, Davis IJ, Argani P, Shukla N, McGill GG, Nagai M, et al. TFE3 fusions activate MET signaling by transcriptional up-regulation, defining another class of tumors as candidates for therapeutic MET inhibition. Cancer Res 2007;67(3):919–29 doi 10.1158/0008-5472.CAN-06-2855.

11. Ho AL, Vasudeva SD, Lae M, Saito T, Barbashina V, Antonescu CR, et al. PDGF receptor alpha is an alternative mediator of rapamycin-induced Akt activation: implications for combination targeted therapy of synovial sarcoma. Cancer Res 2012;72(17):4515–25 doi 10.1158/0008-5472.CAN-12-1319.

12. Pappo AS, Patel SR, Crowley J, Reinke DK, Kuenkele KP, Chawla SP, et al. R1507, a monoclonal antibody to the insulin-like growth factor 1 receptor, in patients with recurrent or refractory Ewing sarcoma family of tumors: results of a phase II Sarcoma Alliance for Research through Collaboration study. J Clin Oncol 2011;29(34):4541–7 doi 10.1200/JCO.2010.34.0000.

13. Olmos D, Postel-Vinay S, Molife LR, Okuno SH, Schuetze SM, Paccagnella ML, et al. Safety, pharmacokinetics, and preliminary activity of the anti-IGF-1R antibody figitumumab (CP-751,871) in patients with sarcoma and Ewing’s sarcoma: a phase 1 expansion cohort study. Lancet Oncol 2010;11(2):129–35 doi 10.1016/S1470-2045(09)70354-7.

14. Sleijfer S, Ray-Coquard I, Papai Z, Le Cesne A, Scurr M, Schoffski P, et al. Pazopanib, a multikinase angiogenesis inhibitor, in patients with relapsed or refractory advanced soft tissue sarcoma: a phase II study from the European organisation for research and treatment of cancer-soft tissue and bone sarcoma group (EORTC study 62043). J Clin Oncol 2009;27(19):3126–32 doi 10.1200/JC0.2008.21.3223.

15. Casanova M, Basso E, Magni C, Bergamaschi L, Chiaravalli S, Carta R, et al. Response to pazopanib in two pediatric patients with pretreated relapsing synovial sarcoma. Tumori 2016:0 doi 10.5301/tj.5000548.

16. Nishio J, Iwasaki H, Ishiguro M, Ohjimi Y, Fujita C, Yanai F, et al. Establishment and characterization of a novel human desmoplastic small round cell tumor cell line, JN-DSRCT-1. Lab Invest 2002;82(9):1175–82.

17. Connell ND, Rheinwald JG. Regulation of the cytoskeleton in mesothelial cells: reversible loss of keratin and increase in vimentin during rapid growth in culture. Cell 1983;34(1):245–53 doi 10.1016/0092-8674(83)90155-1.

18. Markides CS, Coil DR, Luong LH, Mendoza J, Kozielski T, Vardeman D, et al. Desmoplastic small round cell tumor (DSRCT) xenografts and tissue culture lines: Establishment and initial characterization. Oncol Lett 2013;5(5):1453–6 doi 10.3892/ol.2013.1265.

19. Rafehi H, Orlowski C, Georgiadis GT, Ververis K, El-Osta A, Karagiannis TC. Clonogenic assay: adherent cells. J Vis Exp 2011(49) doi 10.3791/2573.

20. Imataka G, Arisaka O. Chromosome analysis using spectral karyotyping (SKY). Cell Biochem Biophys 2012;62(1):13–7 doi 10.1007/s12013-011-9285-2.

21. Schrock E, du Manoir S, Veldman T, Schoell B, Wienberg J, Ferguson-Smith MA, et al. Multicolor spectral karyotyping of human chromosomes. Science 1996;273(5274):494–7.

22. Cheng DT, Mitchell TN, Zehir A, Shah RH, Benayed R, Syed A, et al. Memorial Sloan Kettering-Integrated Mutation Profiling of Actionable Cancer Targets (MSK-IMPACT): A Hybridization Capture-Based Next-Generation Sequencing Clinical Assay for Solid Tumor Molecular Oncology. J Mol Diagn 2015;17(3):251–64 doi 10.1016/j.jmoldx.2014.12.006.

23. Somwar R, Shum D, Djaballah H, Varmus H. Identification and preliminary characterization of novel small molecules that inhibit growth of human lung adenocarcinoma cells. J Biomol Screen 2009;14(10):1176–84 doi 10.1177/1087057109350919.

24. Vojnic M, Kubota D, Kurzatkowski C, Offin M, Suzawa K, Benayed R, et al. Acquired BRAF rearrangements induce secondary resistance to EGFR therapy in EGFR-mutated lung cancers. J Thorac Oncol 2019 doi 10.1016/j.jtho.2018.12.038.

25. Liu Z, Wei P, Yang Y, Cui W, Cao B, Tan C, et al. BATF2 Deficiency Promotes Progression in Human Colorectal Cancer via Activation of HGF/MET Signaling: A Potential Rationale for Combining MET Inhibitors with IFNs. Clin Cancer Res 2015;21(7):1752–63 doi 10.1158/1078-0432.CCR-14-1564.

26. Mattar M, McCarthy CR, Kulick AR, Qeriqi B, Guzman S, de Stanchina E. Establishing and Maintaining an Extensive Library of Patient-Derived Xenograft Models. Front Oncol 2018;8:19 doi 10.3389/fonc.2018.00019.

27. Filion C, Motoi T, Olshen AB, Lae M, Emnett RJ, Gutmann DH, et al. The EWSR1/NR4A3 fusion protein of extraskeletal myxoid chondrosarcoma activates the PPARG nuclear receptor gene. J Pathol 2009;217(1):83–93 doi 10.1002/path.2445.

28. Ritchie ME, Phipson B, Wu D, Hu Y, Law CW, Shi W, et al. limma powers differential expression analyses for RNA-sequencing and microarray studies. Nucleic Acids Res 2015;43(7):e47 doi 10.1093/nar/gkv007.

29. Hanzelmann S, Castelo R, Guinney J. GSVA: gene set variation analysis for microarray and RNA-seq data. BMC Bioinformatics 2013;14:7 doi 10.1186/1471-2105-14-7.

30. Yu G, Wang LG, Han Y, He QY. clusterProfiler: an R package for comparing biological themes among gene clusters. OMICS 2012;16(5):284–7 doi 10.1089/omi.2011.0118.

31. van der Maateen L, Hinton G. Visualizing data using t-SNE. Journal of Machine Learning Research 2008;9:2579–605.

32. Roskoski R, Jr. The ErbB/HER receptor protein-tyrosine kinases and cancer. Biochem Biophys Res Commun 2004;319(1):1–11 doi 10.1016/j.bbrc.2004.04.150.

33. Liu J, Nau MM, Yeh JC, Allegra CJ, Chu E, Wright JJ. Molecular heterogeneity and function of EWS-WT1 fusion transcripts in desmoplastic small round cell tumors. Clin Cancer Res 2000;6(9):3522–9.

34. La Starza R, Barba G, Nofrini V, Pierini T, Pierini V, Marcomigni L, et al. Multiple EWSR1-WT1 and WT1-EWSR1 copies in two cases of desmoplastic round cell tumor. Cancer Genet 2013;206(11):387–92 doi 10.1016/j.cancergen.2013.10.005.

35. Hingorani P, Dinu V, Zhang X, Lei H, Shern JF, Park J, et al. Transcriptome analysis of desmoplastic small round cell tumors identifies actionable therapeutic targets: a report from the Children’s Oncology Group. Sci Rep 2020;10(1):12318 doi 10.1038/s41598-020-69015-w.

36. Tirode F, Surdez D, Ma X, Parker M, Le Deley MC, Bahrami A, et al. Genomic landscape of Ewing sarcoma defines an aggressive subtype with co-association of STAG2 and TP53 mutations. Cancer Discov 2014;4(11):1342–53 doi 10.1158/2159-8290.CD-14-0622.

37. El Beaino M, Liu J, Wasylishen AR, Pourebrahim R, Migut A, Bessellieu BJ, et al. Loss of Stag2 cooperates with EWS-FLI1 to transform murine Mesenchymal stem cells. BMC Cancer 2020;20(1):3 doi 10.1186/s12885-019-6465-8.

38. Saito T, Yokotsuka M, Motoi T, Iwasaki H, Nagao T, Ladanyi M, et al. EWS-WT1 chimeric protein in desmoplastic small round cell tumor is a potent transactivator of FGFR4. Journal of Cancer Science and Therapy 2012;4(10):335–40.

39. Lee SB, Kolquist KA, Nichols K, Englert C, Maheswaran S, Ladanyi M, et al. The EWS-WT1 translocation product induces PDGFA in desmoplastic small round-cell tumour. Nat Genet 1997;17(3):309–13.

40. Werner H, Idelman G, Rubinstein M, Pattee P, Nagalla SR, Roberts CT, Jr. A novel EWS-WT1 gene fusion product in desmoplastic small round cell tumor is a potent transactivator of the insulin-like growth factor-I receptor (IGF-IR) gene. Cancer Lett 2007;247(1):84–90 doi 10.1016/j.canlet.2006.03.027.

